# Force Recruitment of Neuromuscular Magnetic Stimulation Predicted Through High-Resolution Anatomical 3D Models

**DOI:** 10.1101/2021.12.25.474143

**Authors:** S.M. Goetz, T. Weyh, J. Kammermann, F. Helling, Z. Li

## Abstract

Neuromuscular magnetic stimulation is a promising tool in neurorehabilitation due to its deeper penetration, notably lower distress, and respectable force levels compared to electrical stimulation. However, this method faces great challenges from a technological perspective. The systematic design of better equipment and the incorporation into modern training setups requires better understanding of the mechanisms and predictive quantitative models of the recruited forces. This article proposes a model for simulating the force recruitment in isometric muscle stimulation of the thigh extensors based on previous theoretical and experimental findings. The model couples a 3D field model for the physics with a parametric recruitment model, which is identified with a mixed-effects design to learn the most likely model based on available experimental data with a wide range of field conditions. This approach intentionally keeps the model as mathematically simple and statistically parsimonious as possible in order to avoid over-fitting. The coupled model is able to accurately predict key phenomena observed so far, such as a threshold shift for different distances between coil and body, the different recruiting performance of various coils with available measurement data in the literature, and the saturation behaviour with its onset amplitude. The presented recruitment model could also be readily incorporated into dynamic models for biomechanics as soon as sufficient experimental data are available for calibration.

## 1 Introduction

Magnetic stimulation is a method for activating neurons noninvasively through electromagnetic induction with strong and brief magnetic pulses. At present, magnetic stimulation focuses nearly exclusively on the brain (1). Administered transcranially, magnetic stimulation can evoke direct effects, such as motor-evoked potentials (2,3) or phosphenes (4), while certain pulse rhythms or patterns can also modulate neural circuits and shift their excitability with respect to endogenous signals (5). However, the development of magnetic stimulation has been strongly related to the periphery; even the first successful experiments were performed on lower motor fibers and not the brain (6). Long straight axons have been used for studying the basics of excitation since then (7–11).

In neuromuscular applications, magnetic stimulation can serve as an almost pain-free alternative to electrical stimulation (12–26). Classical rehabilitation can be supported by evoking muscle contraction or performing more complex tasks, such as cycling (27). In rehabilitation, neuromuscular stimulation serves to counteract muscle atrophy and to support relearning of movement sequences. Furthermore, orthodromic signals travling from the periphery back to the central nervous system seem to trigger supportive neuroplastic effects (28).

Early neuromuscular magnetic stimulation approaches stimulated the major nerve trunks before they enter the muscle in the hope of achieving strong muscle contraction. The evoked forces are reasonable, but the handling is substantially more complicated than in electrical stimulation and requires experienced operators as the operator has to locate the nerve trunk and place a focal coil very accurately, which is further hampered by contraction and associated shifts in the anatomy. Thus, recent research and clinical efforts prefer the stimulation of the intramuscular axon tree instead, where the procedure is more practical (14, 18, 19, 25). Whereas the activation used to be weaker initially, appropriate coil designs could improve recruitment and demonstrate that better technologies can overcome such weakness and surpass electrical stimulation (17).

However, the design of novel technology for neuromuscular stimulation, including better coil geometries, requires a quantitative understanding of the recruitment. Currently, there is a major knowledge gap between the physics and the neurophysiology of neuromuscular stimulation. Consequently, it is also not clear which physical quantity is responsible for the neuromuscular activation. Thus, also the optimization of coils is rather ad-hoc. The technology for neuromuscular stimulation is therefore only improving slowly and falling behind the rapid developments in transcranial magnetic stimulation (29, 30).

For the brain, in contrast, recruitment of magnetic and electrical stimulation has been studied intensively (31-34) so that both experimental data sets and realistic recruitment models are available (35, 36). Furthermore, such measurements allowed matching models with experimental data so that the dominant physical quantities could be identified (37–39). Although the physical and neurophysiological conditions in the brain are obviously rather different to a peripheral muscle, the work on brain stimulation can still serve for inspiration.

Recent three-dimensional field modeling techniques and experiments provide the basis to study the activation problem quantitatively for the quadriceps femoris muscle and identify the relevant relationships between the field characteristics of the stimulation coil and the muscle activation (41, 43). The experimental study tested a variety of coils that generate widely different field conditions in the thigh and generated further variations through different coil–leg distances with flexible spacers. The wide range of field conditions generated by the combination of both allowed to rule out that the gradient of the electric field plays a substantial role in the stimulation of the intramuscular axon tree, which pervades the muscle densely with fine branchlets, contacts each individual muscle fiber, winding with rather small curvature around them, and forms a high number of terminals.

However, all available analyses in the literature are mere correlation studies, which neither estimate muscle recruitment nor force generation. The available experimental data in the literature, on the other hand, form a sufficiently large data source with enough parameter variation to support the design of a digital twin. The combination of a realistic 3D anatomy model with a parametric recruitment model that estimates the force from anatomical (muscle and fiber anatomy) and physical (induced electric field distribution) output of the 3D model promises to close the gap between the physics and the force response. The data furthermore can serve for the calibration of free parameters to close the present gaps in the understanding.

The presented model estimates the recruitment behavior, which can be observed in isometric stimulation experiments. Appropriate mechanical descriptions are well known in the literature. Riener et al., for instance, propose a sophisticated implementation of Hill dynamics together with biomechanics and a first-order fatigue/recovery description (44). However, the neuromuscular recruitment in such models is usually represented by a sigmoid fitting curve. This work will demonstrate that a realistic recruitment by neuromuscular magnetic stimulation can be estimated directly based on the field conditions to replace ad-hoc sigmoidal fitting curves.

## 2 Model

### 2.1 Anatomy

A high-resolution model of the human thigh was prepared based on the visible-human data set of the US NIH (45). The data include 70mm color photographs of cryosections with 1mm spacing in z-direction, which provide substantially higher resolution than tomography scans and allow a more detailed identification of tissues and particularly boundaries in between. Similar to other models in magnetic stimulation (46–48), the geometry consists of macroscopic regions with dedicated electrical properties and neglects microscopic structures, such as cell membranes. This common approach assures computability. The different classes of segmented tissue elements types include skin, fat, eleven muscles or muscle groups, the femur, blood vessels, and major extramuscular nerve branches, although the latter are not the stimulation target themselves.

The data were segmented with standard image processing methods. The femur and the muscles were identified by simultaneous three-channel analysis of the color data and segmented by thresholding with a manually fine-tuned tolerance band which was chosen accordingly. Furthermore, visible structures that delimit and adjoin the muscular tissue, such as tendons, supported the separation of different muscles along their boundaries and a reconstruction of their surface. The separation of the fat tissue was performed in two stages. A basic frame was obtained from thresholding. As is common in image segmentation, the threshold was determined in the corresponding histograms as a compromise between wrongly identifying other tissue types (false positive) and forming holes due to unclassified regions (false negative). Afterwards, the dataset was cleaned by eroding and reconstructing the mask in order to eliminate image noise and sharp edges.

The data did not exhibit enough contrast at its boundaries for extracting the skin geometry from the images because the embedding gelatin diffused into the skin. As a remedy, this cover was generated artificially by spanning a thin layer of tissue on top of the virtual body which follows the shape of the surface (see Fig. 1 in the middle): The adipose layer was dilated with a three-dimensional Gaussian filter; thresholding that dataset and subtracting all other segmented tissue types as well as still unclassified interior parts formed an approximately 2mm surface layer.

**Figure 1:**
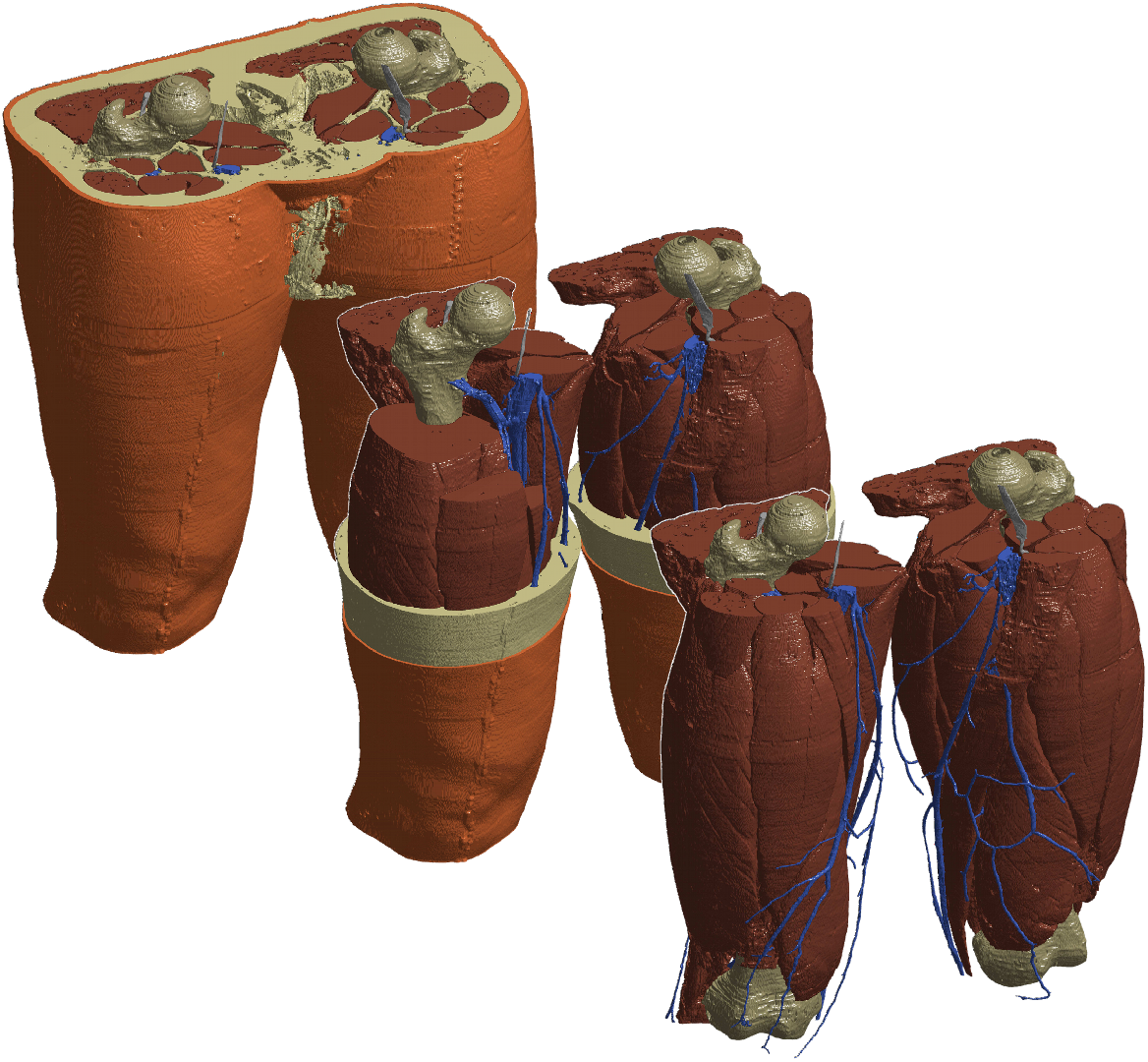
Segmented model in the resolution which was used in the simulation (in the back). In the middle image and in the front, the model is peeled in order to uncover the otherwise hidden structures.

The basis for the blood vessels was prepared by subtracting all already classified regions from the original raw data. From the remaining tissue, small not segmented isles which did not belong to any blood vessels were eliminated by a region-growing algorithm to identify such geometrically isolated subspaces and subsequent removal of unconnected spots. Remaining artifacts and noise were cancelled by three-channel thresholding. Interrupted connections of the grid formed by the vessels were reconstructed by cubic interpolation between the unintentionally separated branches. The major nerve branches (femoral and sciatic nerves as well as the tibial offsprings) were segmented manually from the imaging data.

The boundaries of the individual regions were smoothened with a three-dimensional Gaussian filter in order to suppress unrealistic artifacts and moiré effects. Remaining gaps were filled with the tissue type of the nearest neighbor. For the simulations, only half of the segmented geometry, namely the right thigh, was meshed. The model is depicted in Figure 1 with its various components.

### 2.2 Coils

We modelled four different coil types, namely a standard circular coil (RND15), the racetrack coil RT-120 (MagVenture, Copenhagen, Denmark), the saddle-shaped design APL from (17, 41), and a figure-of-eight coil (MC-B70, MagVenture, Copenhagen, Denmark). The former three devices are taken from the experimental study of (41). The experimental performance of the figure-of-eight coil for magnetic stimulation of the intramuscular nerve structures is not represented in the literature, but was added to predict its force potential using the model calibration to the other coils. The wiring of the coils was extracted from X-ray images and modeled with each individual turn (49, 50). A simplification of the coils to single-turn representations as common in the literature was avoided here (see Fig. 5).

The smaller coils (RND15, RT-120, and MC-B70) were placed with their cross-hairs at the very same location in the center of the proximal third of the right thigh, which is known to evoke the strongest responses in neuromuscular stimulation experiments for the quadriceps muscle group (17). The larger APL coil covers almost the full thigh. The upper edge of the coil ends at the groin. All coils are laterally rotated by 5° in outward direction. The positions are visualized in Fig. 5.

### 2.3 Physics

Due to the low back-action of the induced currents in the tissue, the coils were implemented as unmeshed wires, which determined the magnetic vector potential **A** through the Biot–Savart forward solution (51). The segmented anatomy was meshed with hexahedra and solved with a quasi-static finite volume method (FVM) with more than 70 million volume cells. The FVM enables a high degree of stability and used an established decoupled formulation for stable simulation of eddy currents as detailed in the appendix (41, 52). The electrical characteristics were assigned according to established data in the literature (53) as reported in Table 1.

**Table 1:** Physical properties of the individual tissue regions

|  | Relative permittivity | Conductivity |
| --- | --- | --- |
| | $\epsilon_r$ | $\sigma / \left( \frac{S}{m} \right)$ |
| Skin | $1.1 \cdot 10^3$ | $2.0 \cdot 10^{-4}$ |
| Fat | $2.8 \cdot 10^3$ | 0.024 |
| Muscle tissue | $5.2 \cdot 10^4$ | 0.34 |
| Blood vessels | $1.7 \cdot 10^4$ | 0.31 |
| Major nerve branches | $5.0 \cdot 10^5$ | 0.03 |
| Bone | $8.4 \cdot 10^2$ | 0.082 |

### 2.4 Force Recruitment Model

To this date, 3D models have concentrated on the physical side and studied the distribution of the electric field because the background of eddy currents is well defined and tangible using Maxwell’s equations. However, for the initially requested reproduction of the experimentally observed effects, a physiologic description is essential.

The presented model translates the physical data from the eddy-current simulation into experimentally accessible quantities, namely the isometric force level.

The muscle recruitment and the force generation were represented by a parametric mixed model that is linked to the anatomy and the local electric field in the individual muscles of the 3D model (Fig. 2). The model was subsequently calibrated to experimental data from the literature. We set up several models with different number of parameters and identified the best-suiting model through Schwarz’ Bayesian information criterion (55).

**Figure 2:**
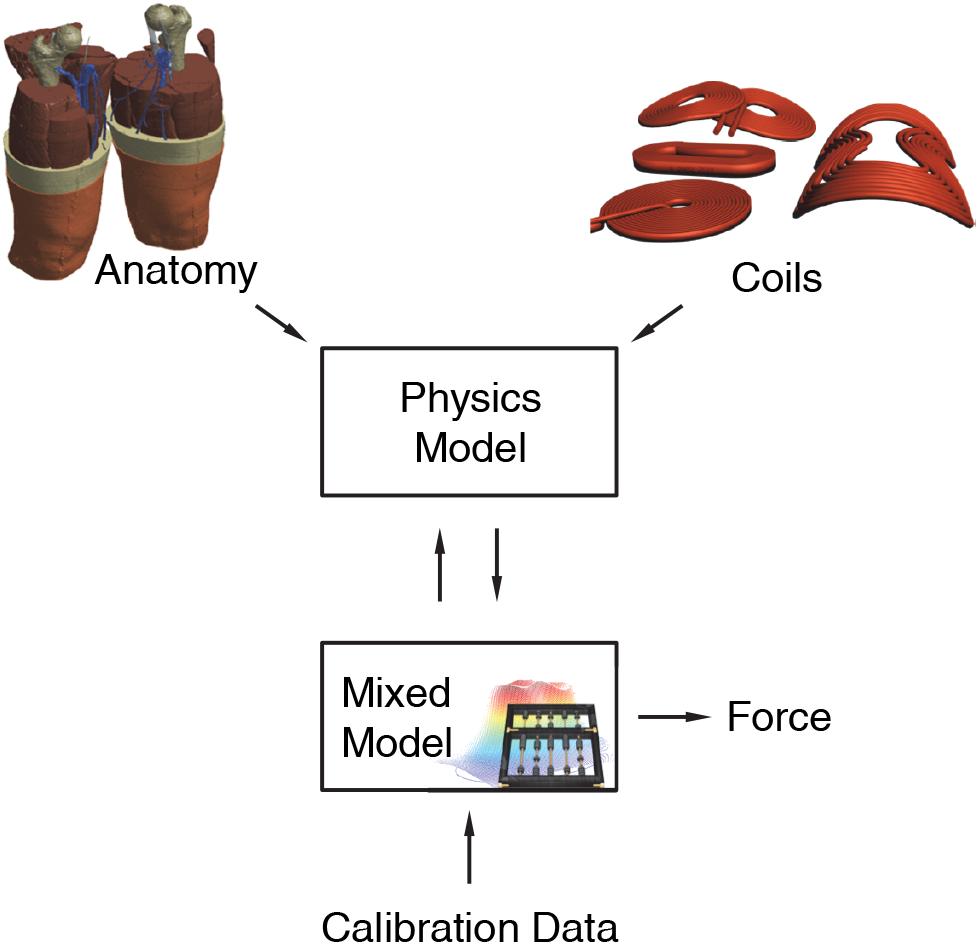
Structure of the overall model with field simulation and parametric mixed recruitment model.

The activation of the intramuscular nerve tree is believed to occur in the fine structure of the axon terminals, close to the neuromuscular junctions (41, 56). For the physiologic approximation here, a microscopic threshold is defined for every junction, which in turn also provides the threshold of the related muscle fiber. Previous research demonstrated that the primary gradient of the electric field is not of greater relevance for explaining the force generation in neuromuscular magnetic stimulation, whereas the electric field strength of the various coils correlates with the response (41, 43). The surface effects at the axon membrane required for an excitation are very effectively generated even in homogeneous fields due to the fine fiber structure with its small curvature radii. This phenomenon appears to reflect the situation in the cerebral cortex, where the activation is also triggered by the field strength rather than by any gradient thereof (57–59). Consequently, we defined a local threshold condition at position **r** within each muscle, which is fulfilled as soon as the norm of the induced electric field **E** exceeds a certain minimum value *E*_th_, ||**E**(**r**)|| < *E*_th_. The free threshold parameter value determines the x-axis of the recruitment curve and the onset of the saturation at higher stimulation amplitudes.

The recruitable force in turn was treated as an individual characteristic. We set up two basic models, one volume-related, one cross-section-area-related as follows.

Most obvious might be estimating the force based on the activated muscle volume, which was used as one model alternative. Due to the fibrous structure of muscles, however, the force generation is not a volume-related issue, nor does the innervation support such an approach (60). Instead, the number of parallel myofibrils determines the peak force during the onset of a contraction in time when no fatigue effects (neither short-term nor endurance effects) are notable, whereas series muscle fibers is widely irrelevant for the tendon force (61–63). This number is approximately proportional to the activated cross-section area perpendicular to the fiber pennation (64, 65).

For extracting the activated share of the physiologic cross-section area of a specific muscle which fulfills the threshold condition acts as an approximation for the force, accordingly. Each muscle of the quadriceps was handled separately in the model. The corresponding cross sections in each muscle were tilted in order to reflect the pennation axis. For the force evaluation, that cross section was taken into account which led to the highest supra-threshold area. The used individual pennation angles with respect to the femur axis were 10° for the m. rectus femoris, 8° for the m. vastus lateralis, −8° for the m. vastus intermedius, and 15° for the m. vastus medialis.

The maximum force level per physiologic muscle cross section is known to be highly individual due to the dependence on the training state, the blood circulation, micro-physiology, the fiber-type composition, and the actual metabolistic conditions (65). For both fundamental model designs, we assumed that the maximum force per area is similar for all members of the relevant extensor muscle group following earlier observations and to keep the model parsimonious (65).

We set up two parametric mixed model alternatives, each with several refinement levels to be calibrated: A first model estimated the isometric force recruitment from the physics model based on the volume activated extensor muscles, i.e., with supra-threshold electric field. The electrical field threshold was a group parameter for all subjects, the force to volume relation, which also determines the maximum recruitable force, individual. As the curvature of the APL coil was too small for some subjects, particularly when further rubber sheet spacers were inserted, a refinement allowed for an individual shift of the APL distance. The second fundamental model used the share of the activated pennation-corrected muscle cross-section. In its simplest form, the threshold field was a group and the force per area an individual parameter. Similarly, in another refinement, the APL coil was given freedom for individual distance shifts.

The force recruitment models were coupled to the 3D physics model through the electric field and the muscle anatomy (see Fig. 2). Each model was trained to experimental data from the literature through mixed-effects maximum likelihood regression (41). We evaluated Schwarz Bayesian information to serve as a model identification criterion that accounts for the different degrees of freedom of the models, particularly in the presence of group and individual parameters.

## 3 Results

The best fitting description used threshold electric field, area-related force, and shift of the APL coil as parameters (log likelihood *L* = −15450); the model identification with Schwarz’ Bayesian information likewise selected this model (*S*_BIC_ = 30919). Second was the models with threshold electric field and area-related force only (*L* = −15675, *S*_BIC_ = 31363); with threshold electric field, volume-related force, and shift of the APL coil (*L* = −15683, *S*_BIC_ = 31385); as well as the one with threshold electric field and volume-related force (*L* = −16261, *S*_BIC_ = 32536).

The electric field threshold across the all data amounted to *E*_th_ = (70.5 ± 21.6) V/m. However, the large spread was caused by only one outlier (see Subject S09 in Fig. 3). This outlier reached 151 V/m, which may be the result of a bad fit of the curved APL coil during the experiments used here and a large effective coil-muscle distance due to a thick subcutaneous adipose layer of the specific subject, which exceeded the one in the model, rather than a really higher local threshold of the intramuscular innervation. The median threshold field was only 65 V/m.

**Figure 3:**
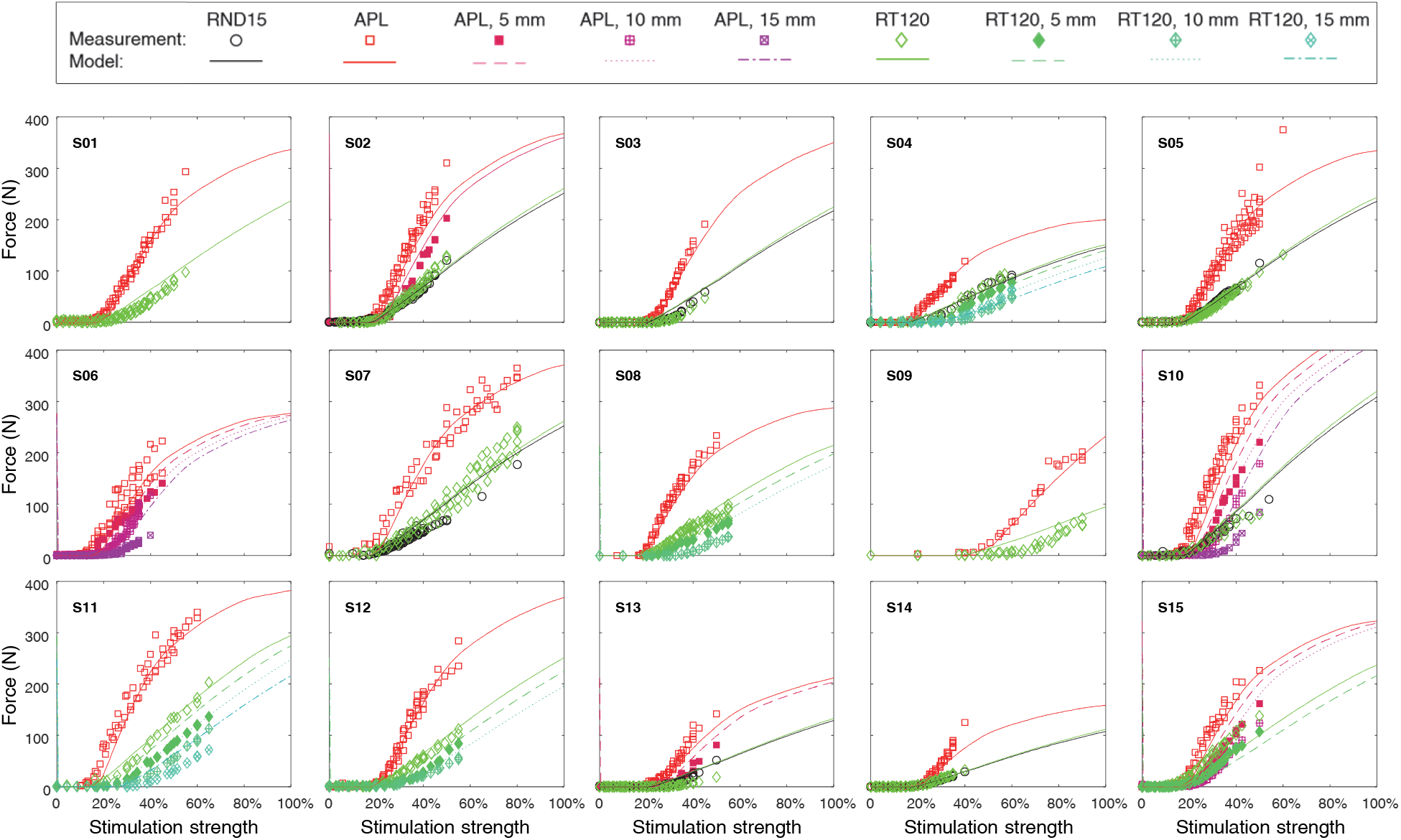
Regression to experimental recruitment data. Each plot displays a different subject with symbols representing the measurement and the lines the force output of the 3D field model in combination with the calibrated mixed model.

Fig. 4 depicts the results of the calibrated model with threshold and maximum force as parameters for the four different coils with various distances from the thigh. Every coil, except the figure-of-eight device, was simulated in its initial condition and for distance values of 5mm, 10mm, and 15mm between the coil and the skin surface in perpendicular direction to reproduce the experimental data. The figure-of-eight coil was incorporated for evaluating its unaltered performance only, since experimental values for the dependence of the distance are not available in the literature. The calibration of the threshold also determines the onset of the saturation for higher amplitudes which agrees well with the experiments and provides validation (see Fig. 4). The simulation reflects the two degrees of freedom, i.e., shift of the recruitment curve with the coil-body distance and slope variations for the different coils, which were observed in experiments (41).

**Figure 4:**
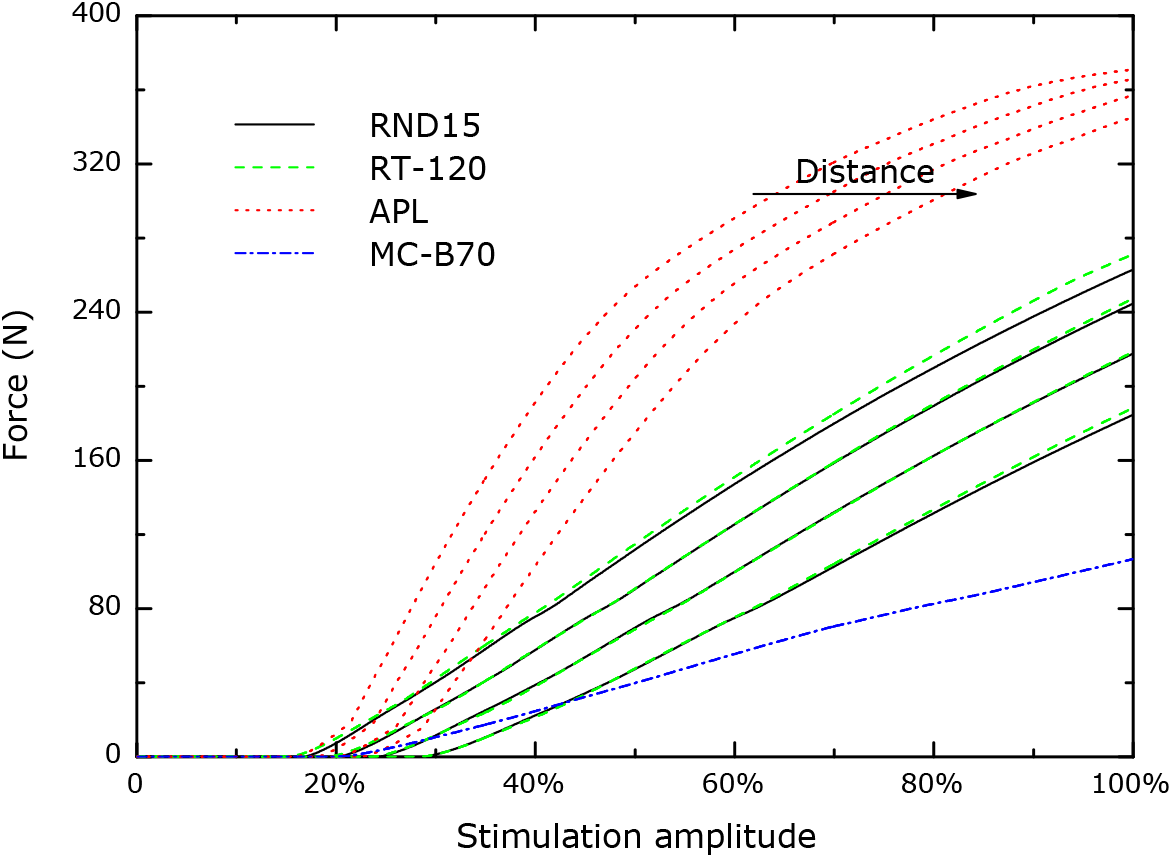
Recruitment curve of all included coils as predicted by the model. Four different distance conditions were simulated (0mm, 5 mm, 10 mm, 15 mm). The slope ratio of the different coils as well as the threshold-shift effect is predicted in good accordance with previous reports.

The distance of the coil from the thigh shifts the threshold in an almost linear way for the observed range. The shift of the recruitment curves in the model is smaller than in experiments, particularly for the APL coil (41). This deviation (29% for the APL coil, 12% for RT-120 in the depicted curves) could be caused by both the experimental setup, in which spacer sheets made of rubber were used. Whereas the coils in direct contact with the thigh ideally match the surface—not least because of the flexibility of the subcutaneous adipose layer—rubber spacers are rather stiff so that the distance increases to a higher extent than the thickness of the sheets. Off-standing edges and air-filled gaps in case of the APL design are difficult to be quantified and simulated correctly. In addition, the simulated model is a standard anatomy and does not represent any individual characteristics of the experimental data which are used here.

The difference in the slopes of the various coils is relatively stable in experiments as well as in the model. The standard circular coil is nearly coincident with the racetrack device RT-120. In the simulations, the APL coil presents an approximately 2.5 times higher slope in the linear range. The figure-of-eight coil—a device which is rarely used for neuromuscular stimulation of the intramuscular axon tree due to low torque but high distress compared to other devices—shows less than half the slope of the standard round coil. The evaluation of the experimental series in (41) led to a ratio of the slopes of the APL and of the standard circular coil of 2.6.

The different performance of the coils can be visualized lucidly by marking the specific subvolume which exceeds the threshold for different points on the recruiting curve. Fig. 5 illustrates the part of the quadriceps muscle (dark) above the threshold field strength. Around the threshold, both coils exhibit just marginal activation. Whereas the increase is rather slow and locally confined for the round coil, the APL device is adjusted to the shape and the size of the quadriceps; the coil even reaches saturation within the output range of the pulse source and for reasonable power. The more homogeneous recruitment might be another, physiologic argument for using coil designs similar to the APL device. The round coil, although frequently used in neuromuscular stimulation, is not able to activate a wider range of fibers, but forms relatively local hot spots. However, these risk conditions could damage relaxed fibers due to the nonphysiological strain acting on them.

**Figure 5:**
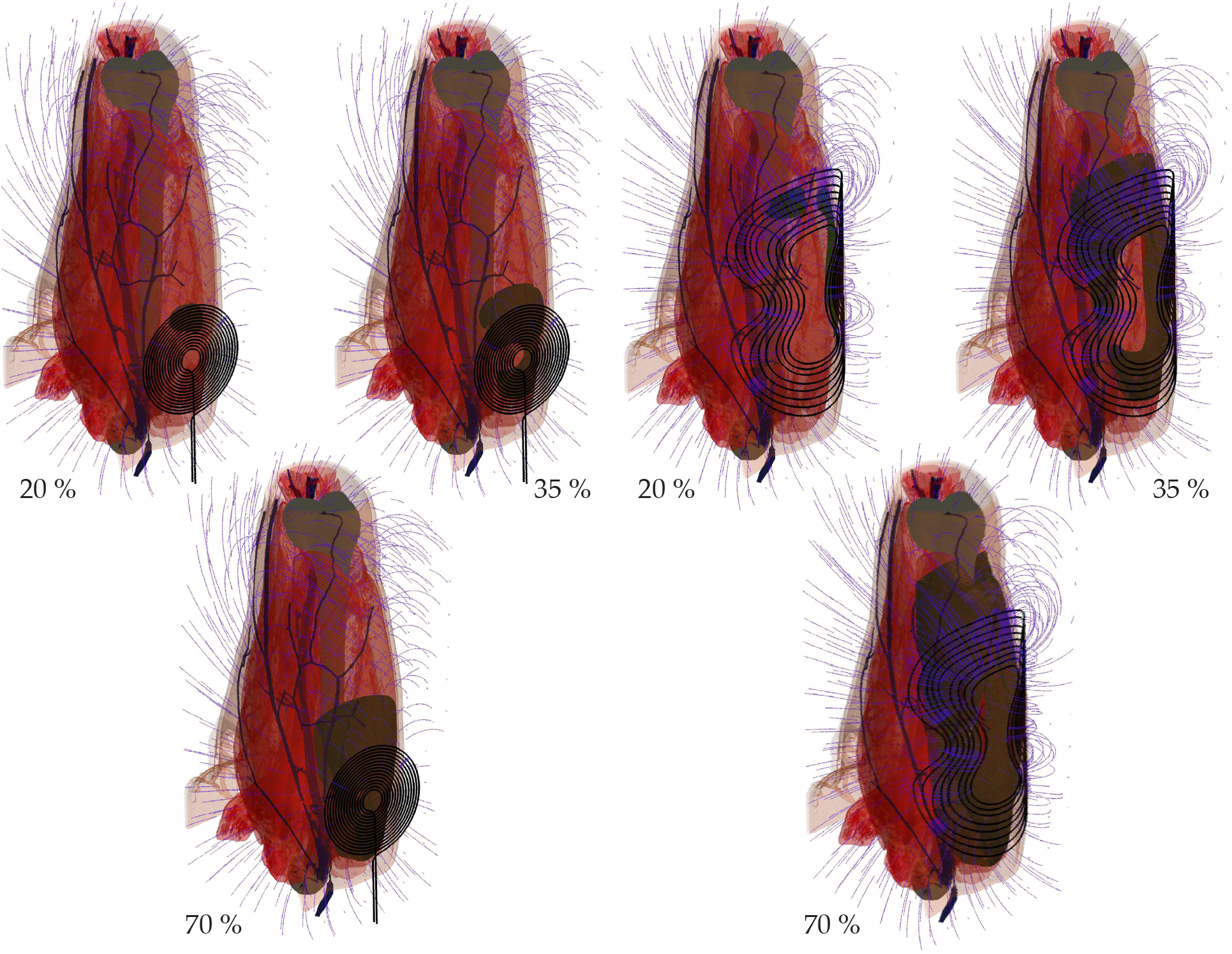
Recruiting for a round circular coil (a) and the APL design (b) at different stimulation amplitudes (20 %, 35 %, and 70% of maximum stimulator output): Whereas the round circular coil shows a rather local activation pattern, the APL device leads to suprathreshold stimulation in almost the whole quadriceps muscle at 70 %. The amount of the muscle volume which fulfils the threshold condition is shaded in dark color in the semitransparent anatomy model; the blue stream lines illustrate the magnetic field or flux.

Although a volume-based approach of the force estimation might be more obvious at first, the growth of the suprathreshold volume in the 3D figures for higher stimulation amplitude is much higher than the experimentally observed force increase. A direct relation of the suprathreshold volume and the force response overestimates the slope of the APL coil (factor of 3.3 instead of 2.6) and predicts a notably lower threshold for this coil compared to the other devices. The difference would correspond to a distance of almost 20 mm which was not observed in the experiments (41).

## 4 Conclusions

Whereas magnetic stimulation has been too weak for daily neuromuscular applications for a long time, more efficient coils eradicated this flaw. A rather simplistic understanding in combination with heuristic trial and error were sufficient for this improvement (17) and allowed the derivation of the coupling factor to estimate the maximum efficiency level (43, 73). This work computationally reproduced the recruiting behavior of neuromuscular magnetic stimulation for the first time. The underlying model is comparably simple, but turned out to be very capable and predicted the different recruiting of various stimulation conditions correctly. If relative torques without absolute force values are sufficient, only one parameter has to be calibrated, i.e., the local intramuscular electrical threshold field. The extraction of this single parameter from measurements entails all remaining effects, such as the experimentally found slope differences of coils and the shift of the recruitment curve for increased coil-thigh distance.

It is very likely that the model can be further simplified, for instance with respect to the anatomic resolution, in order to provide a handy tool for coil designers. In this context, it could support the urgent need of experimentalists and clinicians for adequate equipment as the performance of a design from a drawing board can be subjected to a first, but rather informative quantitative evaluation without large efforts.

Furthermore, the approach was intentionally kept as plain as possible in order to demonstrate that neuromuscular magnetic stimulation behaves macroscopically, i.e., on average notably simpler than the complexity of the microscopic neuromuscular structure might suggest. The parsimony of the model avoids a high number of parameters and factors that might facilitate over-fitting. Still, the model concurs with previous experiments and stimulation studies (14, 74). The simplicity also reflects the need of coil designers in academia and industry for a flexible and usable model.

The model design and calibration became possible due to experimental data of recruitment curves under a sufficient variety of electromagnetic conditions (41). The underlying study evaluated a series of coils and identified two degrees of freedom for changing the magnetic (and induced electric) field conditions across individuals. The essential parameters of the physiologic description—given by the average threshold of the intramuscular axon microstructure and the force per cross section—can be calibrated using measurements. All further characteristics of the recruitment curve, such as the onset of saturation, result from it. In addition, the calibration provides a value for the local electric field strength at the threshold of 65 V/m in the microstructure after excluding one outlier, which is closely related to the microanatomic conditions of the intramuscular axon tree in the neighborhood of the nerve terminals. Interestingly, the threshold value is comparable to corresponding values reported for magnetic stimulation in the brain (66). The model is able to mimic the properties of known muscle stimulation coils quantitatively. It allows predicting the different slopes of various coils correctly and also describes the shift of the threshold with increasing spacing between coil and thigh.

In addition to serving as a tool to design and test new neuromuscular stimulation equipment *in silico*, the model can replace the usually predefined sigmoid functions for the recruiting curve or the behavior of another contractile element in biomechanical models (44, 67, 68). The incorporation into such a framework furthermore provides all temporal aspects of force generation, such as force onset, fatigue and pulse repetition rate, and enables the incorporation into a control loop for functional magnetic stimulation (69, 70). However, this step will require additional experimental data and induces further work in this field.

The quality of the underlying anatomical and geometrical model is relatively high and among the most-detailed in magnetic stimulation. However, even under isometric conditions, muscles change their shape during contraction. For changing knee angles, the situation becomes even more complex. The presented model neglects that. On the one hand, this is a source of error for an accurate reproduction of the physiologic conditions. On the other hand, it keeps the model simple, and the predictive power of the model is sufficiently high after all.

### A FVM Formulation

The formulation used here ans established previously is derived from applying the current continuity to Ampère’s circuital law and introducing the electric field **E** through Ohm’s law. After enforcing Gauß’ law with Poincaré’s lemma, resp. Helmholtz’s theorem through magnetic vector potential with **B** = **curl A** for the magnetic flux density **B** and Coulomb gauging, the governing equations follow

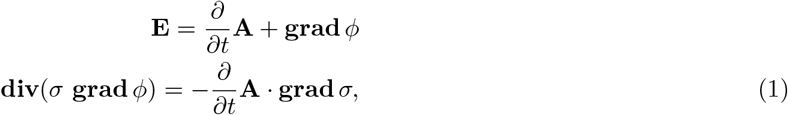

with the local electrical conductivity *σ* and the electrical potential *ϕ*. The FVM turns the differential equation into

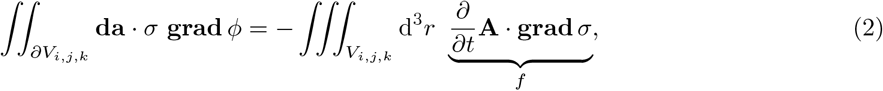

with the volume *V_i,j,k_* of hexahedral volume cell (*i,j, k*). Spatial discretization and first-order finitization of the integrals delivers

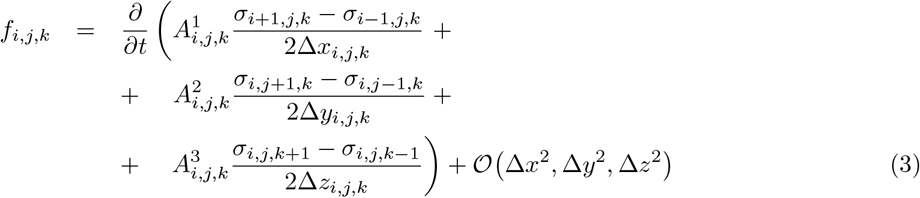

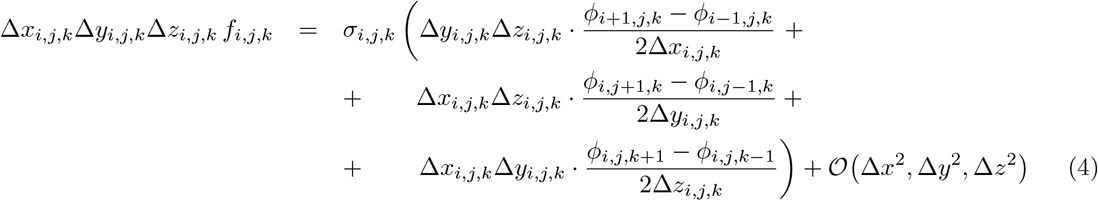

for local variables with dimensions in the superscript. The vector potential for the excitation term *f_i,j,k_* of each cell was provided by the coil’s Biot-Savart solution. The second equation was amended by natural Neumann boundary conditions for *ϕ* on the surface and assembled into a matrix-vector equation (52). Equations were solved by the Gauß-Seidel method.

This formalism can easily be expanded to higher order, also just locally (76):

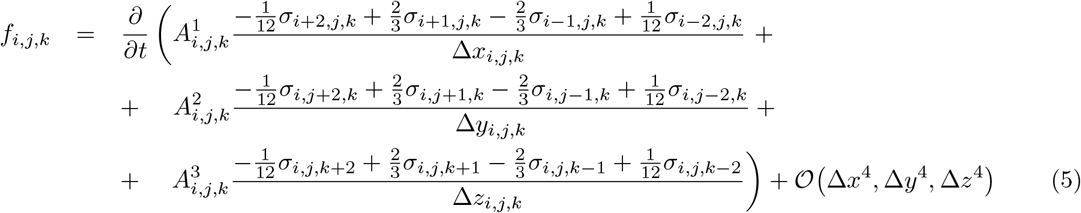

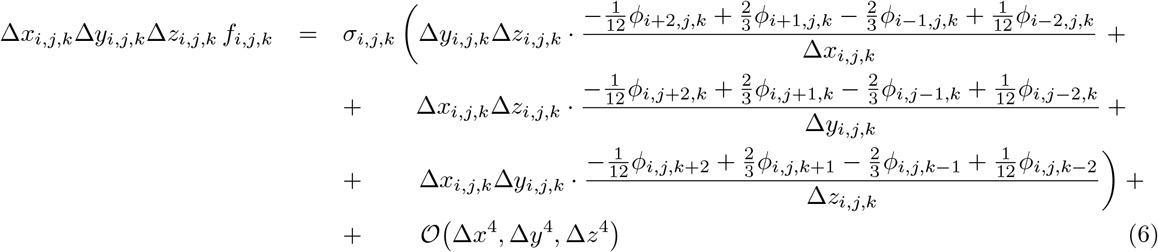

